# Viscoelasticity of globular protein-based biomolecular condensates

**DOI:** 10.1101/2023.12.19.572442

**Authors:** Rachel S. Fisher, Allie C. Obermeyer

## Abstract

The phase separation of biomolecules into biomolecular condensates has emerged as a ubiquitous cellular process. Understanding how intrinsically disordered protein sequence controls condensate formation and material properties has provided fundamental biological insights and led to the development of functional synthetic condensates. While these studies provide a valuable framework to understand subcellular organization via phase separation they have largely ignored the presence of folded domains and their impact on condensate properties. We set out to determine how the distribution of sticker interactions across a globular protein contributes to rheological properties of condensates and to what extent globular protein-containing condensates differ from those formed from two disordered components. We designed three variants of green fluorescent protein with different charge patterning and used dynamic light scattering microrheology to measure the viscoelastic spectrum of coacervates formed with poly-lysine over a timescale of 10^-6^ to 10 seconds, elucidating the response of protein condensates in this range for the first time. We further showed that the phase behavior and rheological characteristics of the condensates varied as a function of both protein charge distribution and polymer/protein ratio, behavior that was distinct to condensates formed with folded domains. Together, this work enhances our fundamental understanding of dynamic condensed biomaterials across biologically relevant length- and time-scales.

## Introduction

Biomolecular condensates, membraneless mesoscale assemblies of proteins and nucleic acids, are increasingly understood to play critical roles in cellular regulation and organization.(1) The formation of these condensed phases was initially shown to be a result of liquid-liquid phase separation (LLPS) of the constituent biomolecules. However there is increasing evidence that biomolecular condensates are complex viscoelastic fluids with formation more readily described by coupled associative and segregative phase transitions (COAST), an umbrella term, that includes LLPS, but more broadly encompasses phase separation coupled to percolation and complex coacervation.(2–6)

The emergent material properties of condensates are important to both the cellular function and their disfunction in the case of disease.(7) Understanding how protein features contribute to the viscoelastic network of biomolecular condensates and consequently dictate emergent properties is therefore key. Multivalent interactions, or transient reversable crosslinks between species, are important drivers of condensate formation.(1, 8, 9) The number of crosslinks, the spacing of crosslinks, and the strength of the molecular interactions forming these transient crosslinks are all key determinants of the underlying viscoelastic network properties.(2, 10, 11) In recent years, intrinsically disordered proteins (IDPs) or disordered regions (IDRs) have emerged as important drivers of condensate formation, as they provide a flexible domain capable of forming a myriad of weak, transient interactions. A major and successful effort has been made to identify sequence level rules that govern the phase behavior of IDPs.(9, 12)

The link between IDP sequence and condensate formation, has been leveraged to successfully design disordered sequences with sticker residues that form functional synthetic condensates.(13) Strategic mutation of endogenous IDPs and design of de novo sequences have both been used to create cellular condensates that can recruit and release cargo.(14) This enables the use of these engineered IDPs to create synthetic compartments that can control enzymatic activity, enhance reaction rates by co-localizing species,(15, 16) or disrupt disease pathways.(17, 18) Synthetic condensates also show great promise in protein delivery with several recent examples of condensates crossing membranes(19) and entering living cells.(20) While IDP’s are important drivers of phase separation, for both endogenous proteins and many of the potential applications of condensates, the phase separating proteins also contain folded domains. In addition to the important biological function, such as catalytic activity, that these globular domains contribute these folded regions can impact phase behavior as well.

Despite the importance of folded domains, the same level of sequence to condensate material property understanding has yet to be established.(21) In the case of a globular protein condensate, the sticky points on the structured domain are not coupled to chain dynamics and cannot move independently. This may result in deviations from the traditional sticky Rouse type understanding that has been used to explain the behavior of IDP condensates. While condensate function may depend on the rheological properties, the biological function, ranging from enzymatic activity to binding specificity, is clearly dependent on the activity of structured domains. Similarly, when designing condensates for applications in protein delivery or metabolic engineering nearly all will incorporate a folded protein. It is therefore crucial to shed light on the role folded domains play in both the physical and material properties of cellular condensates.

With this work we asked the question, how do globular domains contribute to condensate material properties? We used model green fluorescent proteins (GFPs), engineered to be negatively charged, thus allowing them to coacervate with poly-lysine. We investigated how charge distributed across a globular domain versus appended as an intrinsically disordered tag impacts phase separation propensity. Using passive video particle tracking and dynamic light scattering (DLS) microrheology we probed the viscoelasticity of these condensates across a timescale of six orders of magnitude. We found globular protein-based condensates to have viscous behavior at long timescales with subtle differences in terminal viscosities, both between proteins and as a function of protein polymer stoichiometry. Compared to a poly-lysine and poly-aspartic acid polymer control, we found condensates that contained globular GFP to be significantly higher viscosity, interestingly, we found this to be solely due to differences in relaxation timescale. At intermediate timescales we observed a rubbery plateau due to formation of transient sticky interactions. We attributed differences in property to the number and distribution of transient crosslinks formed. At short timescales we found behavior was governed by Zimm type scaling, indicative of the importance of hydrodynamic interactions at short times in the water swollen condensed phases. This work offers fundamental understanding of how globular domains contribute to protein condensates, and the viscoelastic behavior of condensates as a whole.

## Results and Discussion

### Distinct phase behavior of globular and tagged GFPs

To investigate the contribution of folded domains to condensate material properties we compared three negatively charged GFP variants (Figure 1, SI Table 1) and their coacervation with a 30-mer of poly-L-lysine (polyK). The proteins each have an overall charge of approximately -12: in the case of iso-GFP this charge is distributed across the surface of the globular domain; for tag12-GFP the globular domain is net neutral and the excess negative charge is entirely localized to a C-terminal disordered, charged tag; for tag6-GFP the negative charge is split between the globular domain and a C-terminal tag (SI Table 1). For comparison, we also evaluated the phase behavior of this same polyK with a 30-mer of poly-L-aspartic acid (polyD).

**Figure 1.**
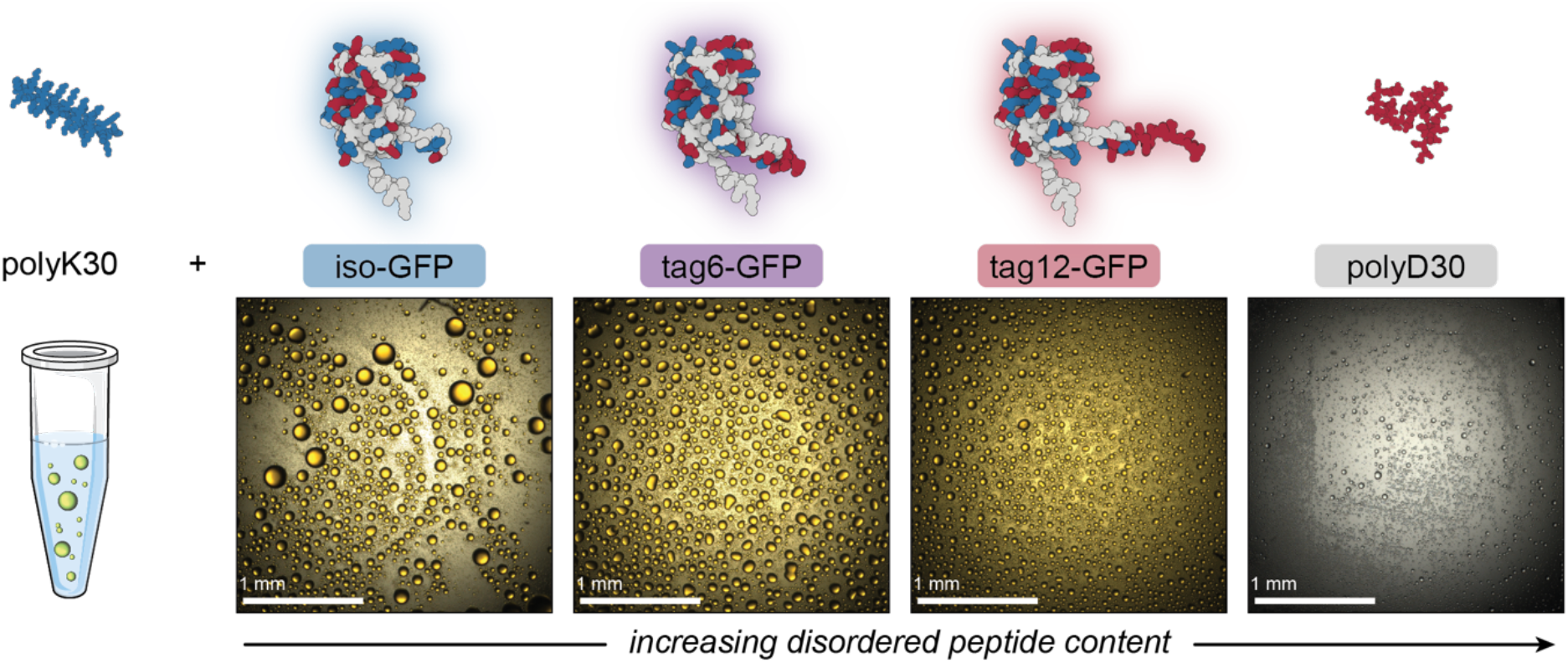
Schematic illustrating the complex coacervates evaluated herein. Coacervates between polyK (DP=30) and three proteins with increasing disordered peptide content were compared to a similar coacervate prepared polyK and polyD (DP=30). The protein structures show negative (red) and positive (blue) residue locations for isoGFP, tag6-GFP and tag12-GFP (left to right). Polypeptide structures were predicted using PEPFOLD3, while protein structures were predicted using AlphaFold. Optical microscopy images show the formation of liquid-like droplets at the optimal mixing ratio for phase separation.

Initially, we explored the phase behavior of iso-, tag6- and tag12-GFP (40 µM) with polyK as a function of mixing ratio and salt concentration. As intrinsically disordered regions are known to promote LLPS(22) and IDP’s with charged residues clustered in blocks as opposed to uniformly distributed have an increased propensity for phase separation we anticipated that the GFP derivatives with disordered tag regions would phase separate more readily. Increasing IDP length is also known to promote phase separation, consequently we expected tag12-GFP to exhibit the highest propensity for phase separation with the cationic peptide.

Phase separation was initially screened by monitoring the turbidity of solutions of the GFPs when mixed with increasing amounts of polyK (Fig 2a). Unlike polyK/polyD coacervates, which had maximum turbidity when the polyions were present at the same ratio (SI Fig 1), all three proteins were found to have maximum turbidity with excess polycation present. This was similarly observed for supercharged proteins including other GFP derivatives and has been attributed to a range of features of globular proteins.(23, 24) This includes the potential for induced charging of ionizable residues upon complexation with polyelectrolytes, the presence of regions of high charge density, or charge patches, as well as the globular structure resulting in local but not global charge neutrality.(23, 24) Interestingly, at low salt conditions maximum turbidity was closer to charge neutral conditions (charge fraction (f^+^) = 0.5 calculated as M^+^/(M^+^+M^-^) where M^+^ and M^-^ are the charge per polymer and protein, respectively) than at higher salt conditions, suggesting a potential role for salt influencing the protein charge state. The largest effect was observed for tag12-GFP indicating the localization of charged residues also played a role, likely because the density of charged residues at the terminus is most prone to induced charging.

**Figure 2.**
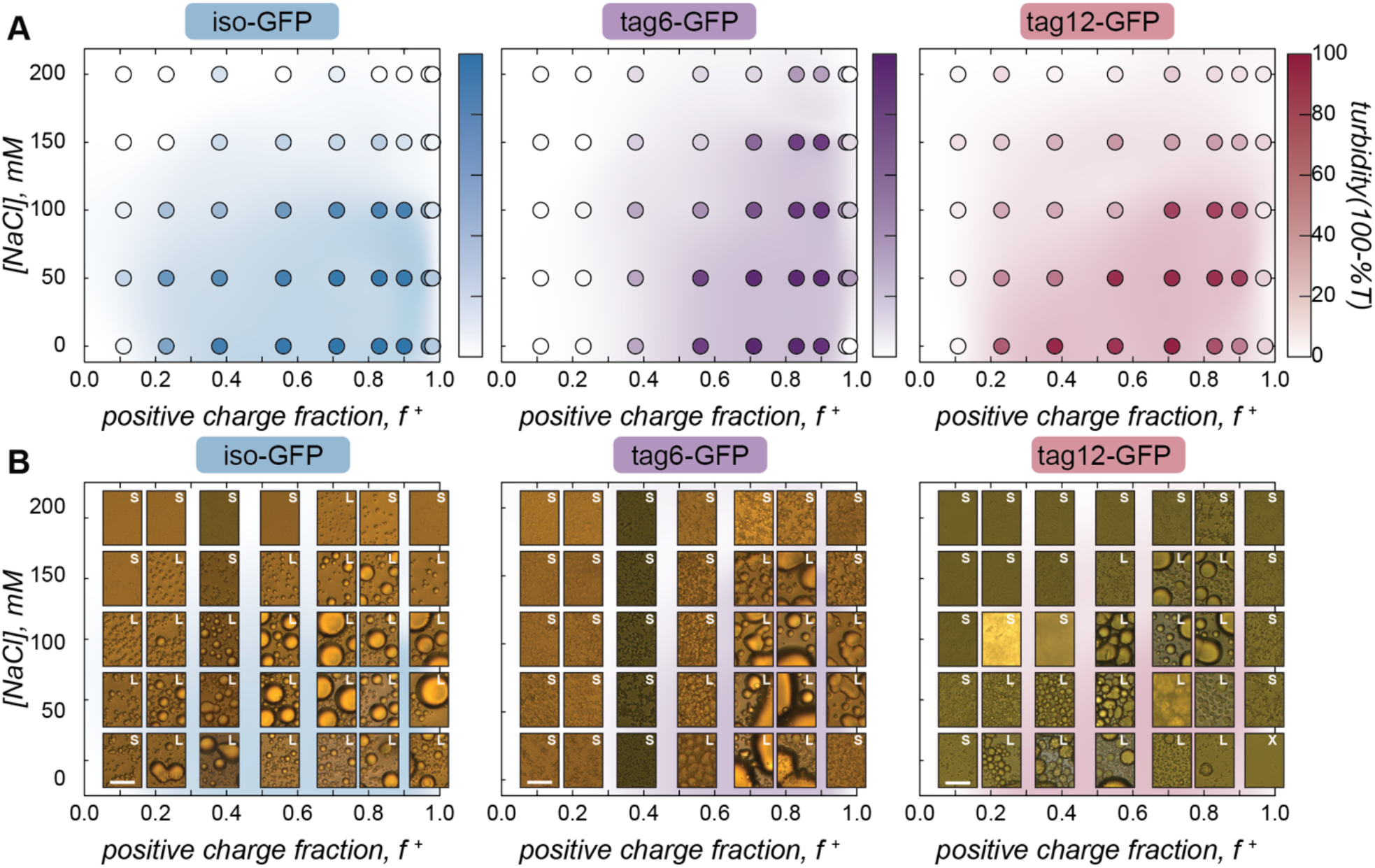
A) iso-GFP, tag6-GFP, and tag12-GFP turbidity as a function of sodium chloride concentration and charge fraction (f^+^) calculated as M^+^/(M^+^+M^-^) where M^+^ and M^-^ are the charge per polymer and protein, respectively. GFP 40 µM, tris (10 mM, pH 7.4). B) Brightfield images of GFP coacervates at varying sodium chloride concentrations and charge fraction. Scale bar 10 µm.

Despite the increased charge blockiness of tag12-GFP, turbidity results suggest it does not have a greater phase separation propensity than iso-GFP (Fig 2a). Upon imaging the samples however, the appearance of the condensed phases was starkly different (Fig 2b). At peak turbidity conditions (f^+^ = 0.8) and salt concentrations below 150 mM NaCl, all three GFPs appear to form spherical liquid-like coacervates, but with increasing amounts of salt tag-GFPs appear to form amorphous aggregates whereas iso-GFP formed liquid-like material. These disparities in the condensed phases formed could result in different degrees of scattering and underrepresentation of solid-like material in turbidity measurements. Therefore, to corroborate maximum coacervate formation at f^+^ = 0.8, protein concentration in the dilute phase was also measured as a function of the mixing ratio of the protein and polymer. Similar behavior was observed, with protein concentration in the dilute phase minimized at this charge ratio (SI Fig 2).

Striking differences between proteins can also be observed when comparing the impact of the positive charge fraction on the material state of the condensed phase. While iso-GFP formed liquid-like coacervates at nearly all conditions tested, tag6-GFP and tag12-GFP formed solid-like material when there was significant excess of either the protein or the polycation (e.g. low and high f^+^) (Fig 2b, SI Fig 3-5). This indicates that addition of a tag potentially increases the binding strength between the GFP and polyK, resulting in the formation of less dynamic condensates or kinetically arrested aggregates.

### Viscosity varies between proteins and as a function of charge ratio

To quantify the effect protein charge distribution has on coacervate material properties, we used video particle tracking microrheology (VPT) to probe the viscosity of the GFP-polyK condensates. Fluorescent microspheres were embedded within condensates (Fig 3a) and widefield microscopy was used to track their trajectories and ultimately report on condensate viscosity.

**Figure 3.**
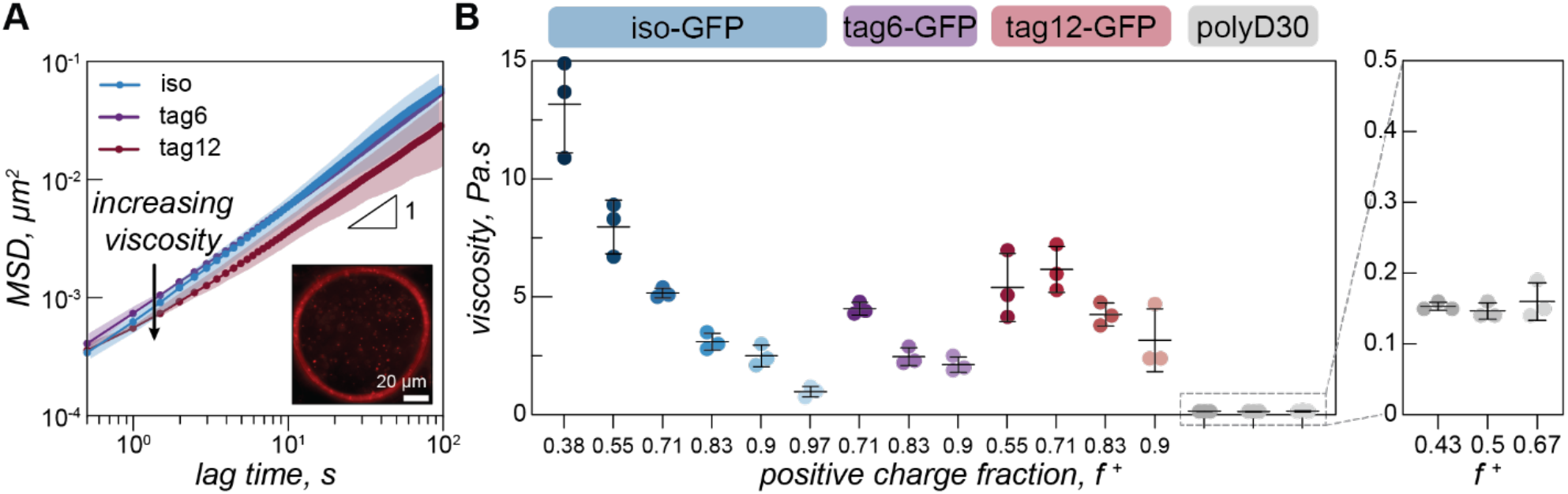
Video particle tracking. A) Mean squared displacement as a function of lag time for iso-GFP (blue), tag6-GFP (purple) and tag12-GFP (maroon) with polyK (f^+^ = 0.7). Plots depict average of three independent measurements. Shaded areas show standard deviation. Inset: Single frame from particle tracking video of iso-GFP-polyK (f^+^ = 0.9) condensate with 500 nm beads embedded. Scale bar is 20 µm. B) Viscosity for iso-GFP (blue), tag6-GFP (purple) and tag12-GFP (maroon) condensates when prepared with 40 µM protein and different polyK concentrations. Iso-GFP coacervates were prepared with 6 polyK concentrations: 10, 20, 40, 80, 148, 500 µM, corresponding to f^+^ = 0.38, 0.55, 0.71, 0.83, 0.9 and 0.97. Similarly, tag6-GFP coacervates were prepared with polyK at 40, 80 and 148 µM and tag12-GFP with polyK at 20, 40, 80 and 148 uM. Three separate measurements (data points), average (black line), and standard deviation (error bars), are shown. Enlarged portion of plot displays viscosity of polyK-polyD coacervates (grey) at three different charge ratios f^+^ = 0.43, 0.5, and 0.67.

Due to the increased charge blockiness in the tagged GFP variants, we hypothesized that condensates containing these GFPs may have higher viscosities. However, we find that viscosities for all three proteins are only subtly different (Figure 3b). At a positive charge ratio of 0.7, tag12-GFP, tag6-GFP, and iso-GFP have a viscosities of 6.2 ± 1.0 Pa.s, 4.5 ± 0.3 Pa.s, and 5.1 ± 0.2 Pa.s respectively. Interestingly it is tag6-GFP, not iso-GFP, that has the lowest apparent viscosity.

Due to the differences between GFP phase diagrams, we hypothesized that viscosity differences could be due to location in phase space. We therefore measured the viscosity as a function of charge ratio. We varied charge ratio by keeping a constant GFP concentration and increasing the polyK concentration. Iso-GFP formed large coacervates amenable to microrheology measurements over the broadest range of conditions, consequently a broader range of concentrations were investigated. We find a general trend of decreasing viscosity with increasing polyK concentration for all three proteins. However, the viscosity of tag12-GFP coacervates appears to plateau or slightly decrease between a charge fraction of 0.55 and 0.7 and the overall change in viscosity across the measured range is less significant than that for iso-GFP.

Interestingly we do not observe similar charge ratio dependent behavior for coacervates formed from polyK/polyD (Fig 3d). Here the viscosity appears to remain constant, an effect previously observed and attributed to one preferred polycation/polyanion ratio within the condensed phase, maintained by changes in the coacervate volume to compensate for initial condition changes.(25, 26) A change in coacervate viscosity can be induced by changing the polycation/polyanion ratio within the dense phase,(27–29) suggesting that unlike in the polyK/polyD coacervates a compositional change in the condensed phase may take place in the GFP coacervates as a function of charge ratio. We also found the viscosity of the polyK/polyD was significantly lower than that of the GFP containing coacervates. This is consistent with previous theoretical and experimental work highlighting the higher density of coacervates containing globular as opposed to linear components.(21, 30)

### Condensates: a transient viscoelastic network

On the timescale of our particle tracking measurements all coacervates behaved as pure viscous fluids. To access the high frequency range required to probe rheological response corresponding to fast segmental chain dynamics and gain more detailed information about the fluid network, we turned to dynamic light scattering microrheology (DLSμR). This technique has previously been shown to reproduce oscillatory macrorheology results for synthetic polymer gels and been used to characterize precious biological fluids such as mucus.(31–33) This approach is advantageous as proteins are liable to denature at elevated temperatures and the coacervate structure and density can vary significantly at elevated salt concentrations, preventing the use of temperature or salt-superpositioning as has been done for synthetic complex coacervates.(25, 34–37)

Comparing the mean squared displacement determined from VPT microrheology to the MSD calculated from DLS for iso-GFP and polyK (40 µM) coacervates (Fig 4a) highlights the different time regimes over which these techniques report. Both cover the 0.5 to ∼10 second time regime, and while there are some differences in the observed absolute value both scale as ∼ 1 and similar trends between samples were observed. While the slope of 1 found from VPT indicated beads were moving in a purely viscous fluid, at timescales faster than 0.5 seconds, the shortest time interval used for VPT, the MSD of the embedded beads measured by DLS can be seen to plateau, indicating non-Brownian behavior. The corresponding complex moduli (Fig 4b) displays three distinct regions: a terminal relaxation or flow region (blue region), a rubbery plateau due to formation of a transient network (white region), and a Rouse-like transition region due to local monomer relaxation (green region). This is qualitatively similar to observations made in superposition experiments with polymer complex coacervates. Each of these regions corresponds to different potential interactions driving the observed viscoelastic behavior, shown in the schematic that represents the dominant interactions occurring within each timescale (Fig 4c).

**Figure 4.**
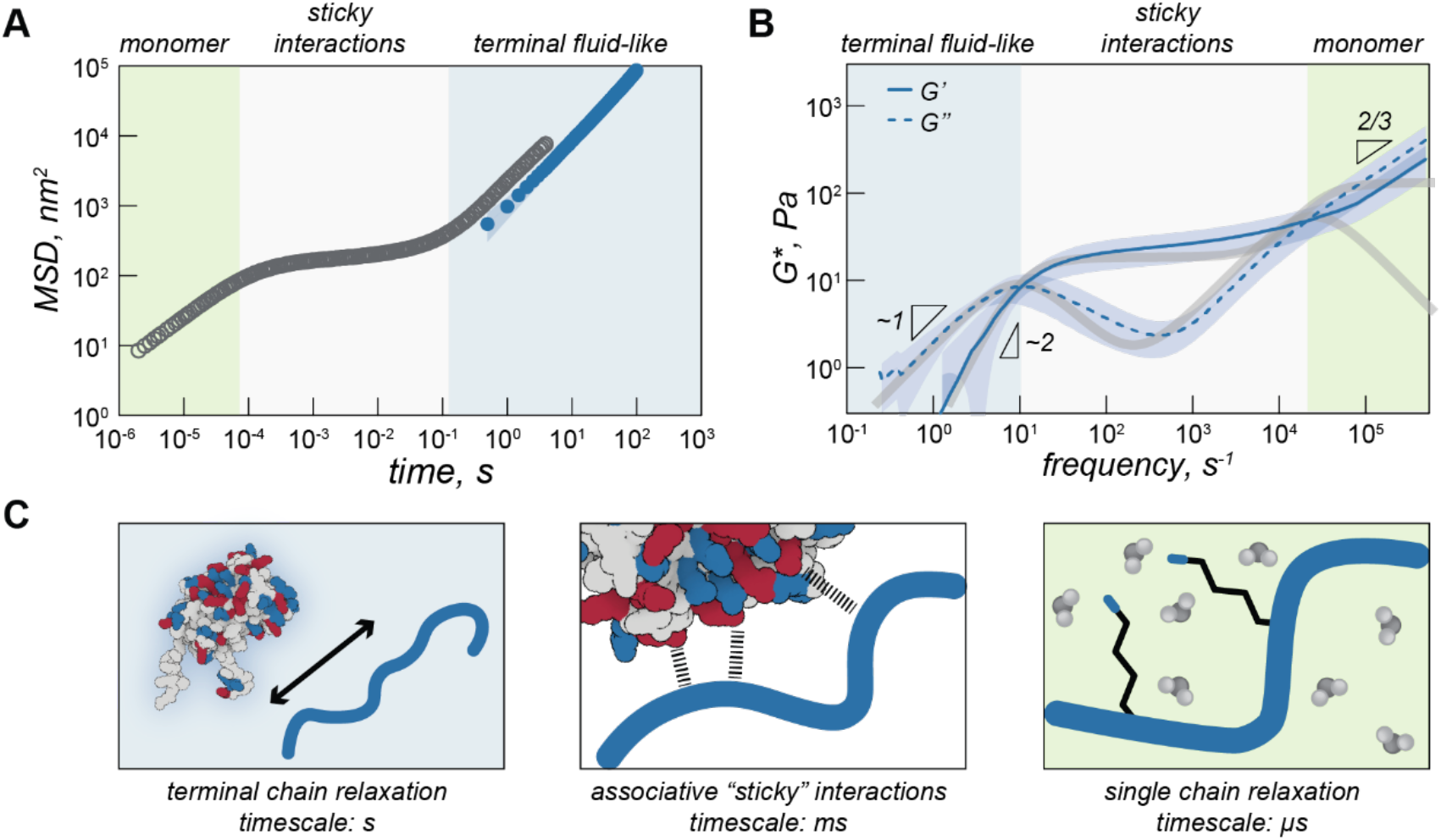
A) Mean square displacement (MSD) of iso-GFP coacervates (polyK 40 µM) determined from DLS microrheology (blue) and MSD determined from video particle tracking microrheology (grey). Background colored regions qualitatively illustrate different timescales. B) Complex modulus of iso-GFP from DLS rheology. G’ solid line, G’’ dashed line. Grey lines represent two-component Maxwell fit. C) Schematic illustrating protein polymer behavior at different timescales. At longest time scales (blue background) proteins and polymers can flow. At intermediate times (white background) behavior is dominated by interactions between charged residues on GFP and polyK. At short timescales (green background) residue interactions with water dominate.

At low frequencies, or the longest timescale probed, condensate behavior is dominated by the loss modulus (G”, indicated with a dashed line) indicating predominantly fluid behavior at long timescales where protein and polymer have time to disassociate. This region is shaded blue in Fig 4a, b & c. In this region, we find that G’ scales as ω^1^ (slope = 1.9) and G’’ scales as ω^2^ (slope = 0.97), which is consistent with the scaling expected for the viscoelastic response of unentangled ideal polymer chains. This is also consistent with the apparent liquid behavior we observed from video particle tracking microrheology as well as with previous work indicating condensates are Maxwell fluids.(3, 5, 26) A two component Maxwell model, with a slow relaxation originating from sticker interactions and a fast relaxation due to Rouse motion of the polymer between sticky bonds, largely described the behavior at long and intermediate time scales but did not fully capture the relaxation behavior at the shortest times scales.

The characteristic relaxation time *r*_*rep*_, the reciprocal of the frequency at which the storage and loss modulus cross, represents the longest time taken for the biopolymers to disassociate from the network and form new crosslinks. For iso-GFP/polyK condensates we can see this occurs at ∼100 ms (Fig 4b, Table 1). Interestingly we observe a much shorter relaxation time, ∼ 3 ms, for the polyK-polyD coacervates (Fig 5f). At shorter timescales than *τ*_*rep*_, we see elastic behavior begin to dominate and a marked plateau in the elastic modulus. This is characteristic of a network held together by sticky interactions, here transient crosslinks formed by electrostatic interactions between GFP and polyK.

**Table 1.**
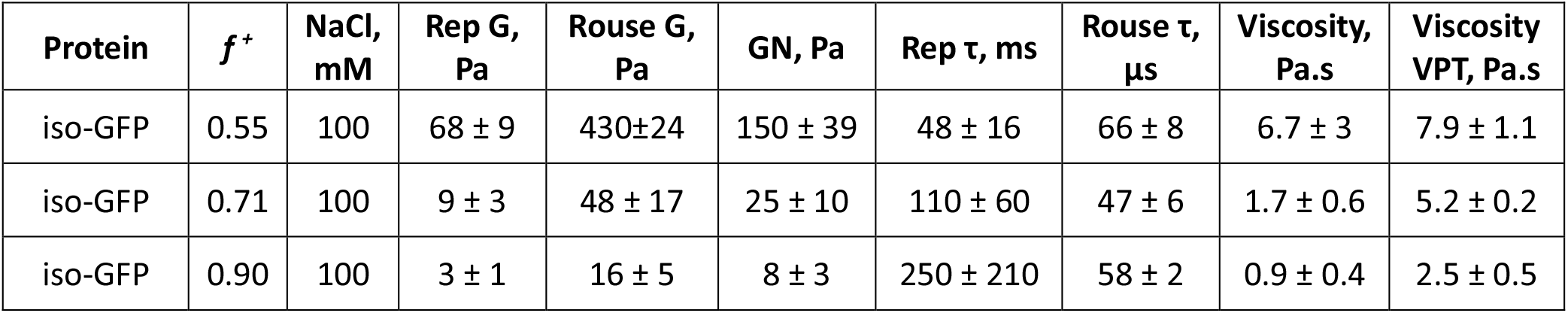
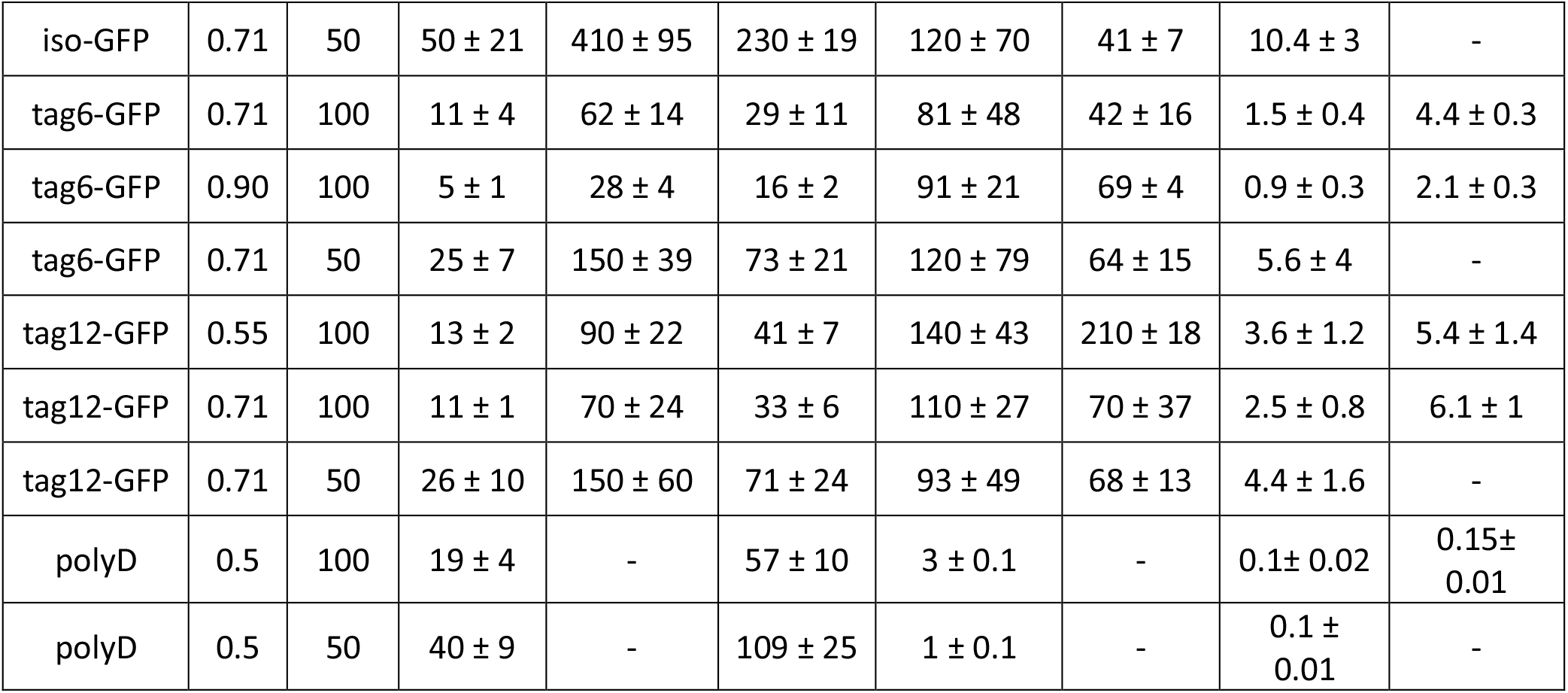
Relaxation times, plateau moduli, and terminal viscosities of condensates as determined by DLS and VPT microrheology.

| Protein | $f^+$ | NaCl, mM | Rep G, Pa | Rouse G, Pa | GN, Pa | Rep $\tau$, ms | Rouse $\tau$, μs | Viscosity, Pa.s | Viscosity VPT, Pa.s |
| --- | --- | --- | --- | --- | --- | --- | --- | --- | --- |
| iso-GFP | 0.55 | 100 | 68 ± 9 | 430±24 | 150 ± 39 | 48 ± 16 | 66 ± 8 | 6.7 ± 3 | 7.9 ± 1.1 |
| iso-GFP | 0.71 | 100 | 9 ± 3 | 48 ± 17 | 25 ± 10 | 110 ± 60 | 47 ± 6 | 1.7 ± 0.6 | 5.2 ± 0.2 |
| iso-GFP | 0.90 | 100 | 3 ± 1 | 16 ± 5 | 8 ± 3 | 250 ± 210 | 58 ± 2 | 0.9 ± 0.4 | 2.5 ± 0.5 |
| iso-GFP | 0.71 | 50 | 50 ± 21 | 410 ± 95 | 230 ± 19 | 120 ± 70 | 41 ± 7 | 10.4 ± 3 | - |
| tag6-GFP | 0.71 | 100 | 11 ± 4 | 62 ± 14 | 29 ± 11 | 81 ± 48 | 42 ± 16 | 1.5 ± 0.4 | 4.4 ± 0.3 |
| tag6-GFP | 0.90 | 100 | 5 ± 1 | 28 ± 4 | 16 ± 2 | 91 ± 21 | 69 ± 4 | 0.9 ± 0.3 | 2.1 ± 0.3 |
| tag6-GFP | 0.71 | 50 | 25 ± 7 | 150 ± 39 | 73 ± 21 | 120 ± 79 | 64 ± 15 | 5.6 ± 4 | - |
| tag12-GFP | 0.55 | 100 | 13 ± 2 | 90 ± 22 | 41 ± 7 | 140 ± 43 | 210 ± 18 | 3.6 ± 1.2 | 5.4 ± 1.4 |
| tag12-GFP | 0.71 | 100 | 11 ± 1 | 70 ± 24 | 33 ± 6 | 110 ± 27 | 70 ± 37 | 2.5 ± 0.8 | 6.1 ± 1 |
| tag12-GFP | 0.71 | 50 | 26 ± 10 | 150 ± 60 | 71 ± 24 | 93 ± 49 | 68 ± 13 | 4.4 ± 1.6 | - |
| polyD | 0.5 | 100 | 19 ± 4 | - | 57 ± 10 | 3 ± 0.1 | - | 0.1 ± 0.02 | 0.15 ± 0.01 |
| polyD | 0.5 | 50 | 40 ± 9 | - | 109 ± 25 | 1 ± 0.1 | - | 0.1 ± 0.01 | - |

**Figure 5.**
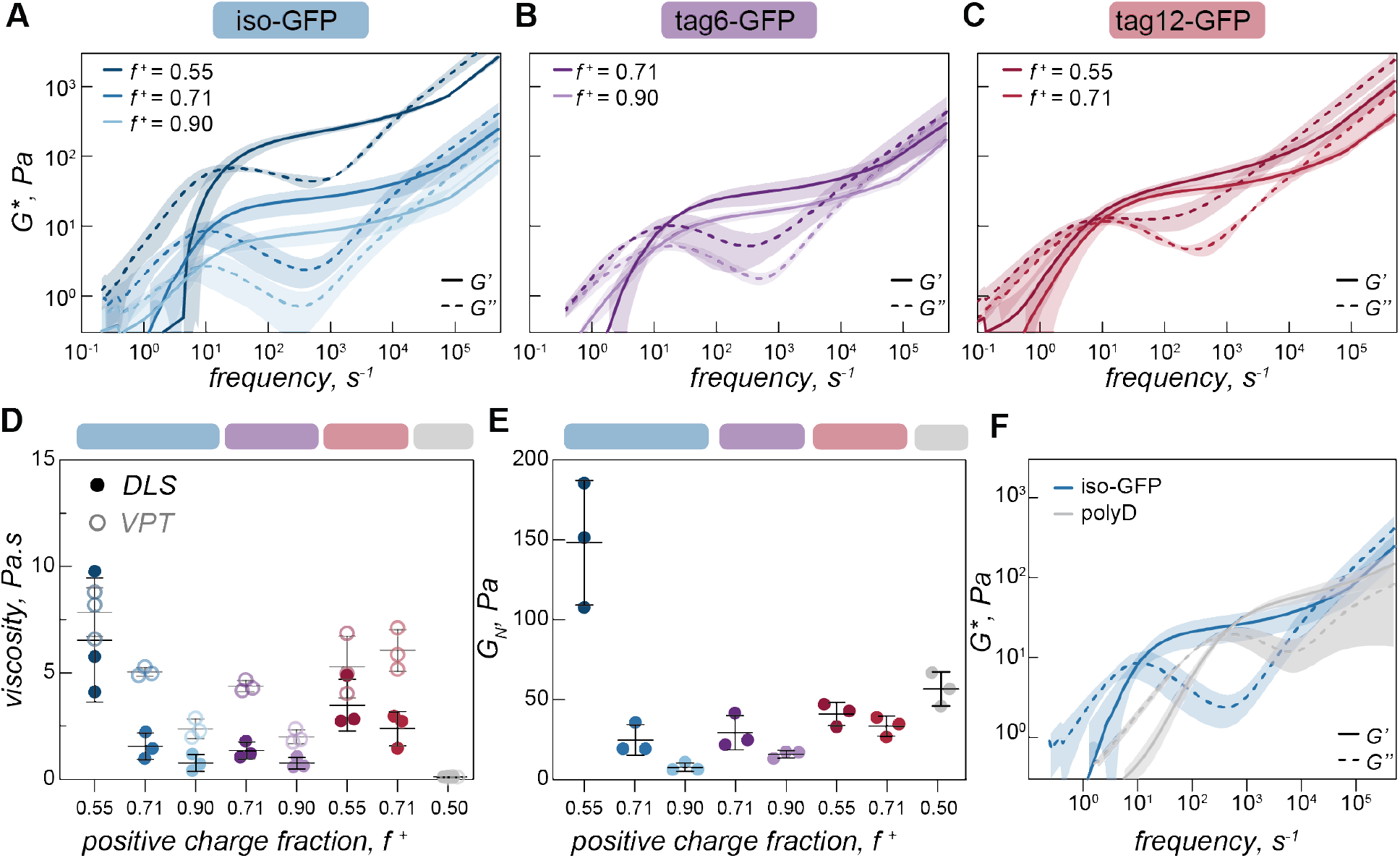
A) Complex modulus of iso-GFP (40 µM) and polyK coacervates at different charge ratios (f ^+^ = 0.55, 0.71, 0.90,) B) tag6-GFP (40 µM) and polyK at different charge ratios (f ^+^ = 0.71, 0.90), C) tag12-GFP (40 µM) and polyK at different charge ratios (f ^+^ = 0.55, 0.71). D) Viscosity (η_0_=2G_0_τ_0_) from DLS (filled circles) and VPT (empty circles) for iso-GFP (40 µM) and polyK coacervates at different charge ratios (f ^+^ = 0.55, 0.71, 0.90) (blue); tag6-GFP (40 µM) and polyK at different charge ratios (f ^+^ = 0.71, 0.90) (purple), tag12-GFP (40 µM) with polyK at different charge ratios (f ^+^ = 0.55, 0.71) (red) and polyD30 (f ^+^ = 0.71) (grey). E) Plateau modulus (G_N_) of iso-GFP (40 µM) and polyK coacervates at charge ratios f ^+^ = 0.55, 0.71, 0.90 (blue), tag12-GFP (40 µM) with polyK at charge ratios f ^+^ = 0.55, 0.71 (purple), tag12-GFP (40 µM) with polyK at charge ratios f ^+^ = 0.55, 0.71 (red) and polyD30 (f ^+^ = 0.71) (grey). F) Complex modulus of iso-GFP (blue), tag6-GFP (purple) and tag12-GFP (red) (40 µM) with polyK (40 µM, f+ = 0.71) at a NaCl concentration of 50 mM. Inset shows plateau modulus (G_N_) at 50 mM NaCl concentration (solid circles) and 100 mM NaCl (open circles).

At the highest frequencies, we find that the elastic modulus scales as ω^2/3^. This region describes interactions occurring at the fastest time scales, typically monomer interactions. Two of the most common methods to describe polymer behavior in this regime are the Rouse and Zimm models.(38) Scaling of G’ as ω^2/3^ in the high frequency range is characteristic of the Zimm model, which differs from a Rouse model (G’ ∼ ω^½^) by asserting the effect of hydrodynamic interactions between polymer monomers and solvent dominate over monomer-monomer interactions, the dominate interaction governing behavior in the Rouse model. A Zimm model describes well the behavior of dilute polymers where long-range interactions like hydrodynamics become increasingly important. This suggests that polyK is in the dilute solvent accessible environment. Due to this, the absence of a second rubbery plateau,(35) and the short length of the polyK (N ≈ 30), these results indicate that there is no polymer chain entanglement. Interestingly as salt is decreased, we see a shift in modulus indicating increased material strength and scaling in the high frequency range as ω^2/3^ appears to slightly decrease (Fig SI 8, Table 1). This suggests that Rouse-like monomer-monomer interactions become increasingly important as the coacervate becomes more dense and less fluid with decreasing solution ionic strength.

#### Protein polymer charge ratio impacts material strength and relaxation time

Having observed changes in terminal viscosity as a function of charge ratio with VPT we wanted to interrogate this effect over a broader time range. Due to the larger volumes required for DLS rheology a subset of charge ratios were measured for the three GFP variants (*f+* = 0.55, 0.71, and 0.90 for iso-GFP (Fig 5a), *f*^*+*^ = 0.71 and 0.90 for tag6-GFP (Fig 5b), and *f+* = 0.55 and 0.71 for tag12-GFP (Fig 5c)). Similar to the trend of decreasing viscosity with increasing polyK concentration observed from VPT, we see a decrease in both zero shear viscosity (Fig 5d) and in the elastic modulus (Fig 5e), indicating decreased material strength and increased fluidity.

These changes in the complex modulus could be due to changes in sticker strength or fewer sticker interactions. If these coacervates were dominated by polymer behavior and governed by a sticky-Rouse type mechanism, a decrease in material strength due to fewer or weaker interactions, would be expected to occur alongside a corresponding increase in the relaxation frequency towards the higher frequency range, indicating that polymers within the condensate can relax more easily. For tag12-GFP this expected shift is observed, albeit subtly. Conversely, for tag6-GFP and iso-GFP there appears to be a small decrease in crossover frequency with decreased network strength. The error associated with the crossover times is large in comparison to subtle changes we observe in the average value between conditions making it unclear if this difference is statistically significant, preventing any conclusive findings of discrepancies with traditional sticky rouse behavior. However, this likely indicates that the proteins are making a significant contribution towards the material properties. We hypothesize that as the polyK concentration increases, the relative proportion of GFP sticky points to polyK molecules decreases, thus increasing fluidity. We hypothesize that this change in ratio also impacts the proportion of sticky interactions that contribute to the network as compared to those that act as dangling ends. Each time a sticky bond breaks, the most likely bond to form is between the same partners, but the characteristic relaxation time should be considered the timescale for a bond to reform with a new partner, not the timescale for bond breaking. The arrangement of neighboring interaction sites, which can be precisely controlled on globular protein surfaces, is key to material properties. For tag12-GFP the localization of charges to one region could increase the probability of one polyK forming multiple interactions with the same GFP, thus not contributing to the network elasticity. This results in a narrower range of conditions where viscoelastic fluids form and an increased propensity towards small aggregated clusters. For iso-GFP, as the charged patches are distributed across different regions, this potentially increases the probability that interactions on one GFP will be with multiple different polyK. This leads to viscoelastic fluids forming over a broad range of conditions. The larger change in modulus observed could be indicative of different charge patches with different strengths. When polyK is the limiting component only dominant GFP charge patches interact whereas when polyK is in excess less charged regions may also become potential sticky points. This could suggest that a spectrum of different clusters form impacting the material response.

#### Polymer-polymer condensates exhibit faster relaxation timescale

From VPT experiments we observed significantly lower viscosity for polyK-polyD, polymer-polymer condensates (Fig 3b) than those containing a globular GFP. Using DLS-rheology we observe a similarly lower terminal viscosity (Fig 5d). Interestingly, we find the plateau modulus to be of a similar magnitude to the globular protein condensates indicating interactions of a similar strength (Fig 5e, 5f). The decrease in viscosity is therefore due to the faster timescale over which the transient crosslinks form and relax, not due to differences in interaction strength (Fig 5f). We attribute this effect to the increased degrees of freedom available to a linear molecule which has segmental flexibility compared to a globular protein which has only translational and rotational degrees of freedom.

## Conclusion

The material properties of condensates are diverse, encompassing dynamic fluids, gel-like solids, glasses and aggregates.(1, 39, 40) Changes in fluidity can impact biological function and transitions from dynamic to more solid like states are associated with pathologic response.(39) Consequently, determining the emergent properties of condensates, such as their viscoelasticity, is of great importance. Condensates are composed of a range of biomolecules, the diffusion of these molecules, the interaction between different species (i.e bond lifetime), and global properties of the condensates as a whole, all occur at different timescales. It is therefore imperative to understand material response at both long and short times, to capture all timescales relevant for function. Here we have determined the viscoelastic spectrum of model GFP and polyK condensates over a broad range of timescales, providing insight into these different regimes.

Particle tracking rheology has been used to characterize a number of *in vitro* protein or peptide-based biomolecular condensate systems.(41) This technique typically allows the MSD between 0.1 to 100 s to be measured. As with the condensates measured here, several display liquid-like behavior over this time regime,(42–44) while for other systems features characteristic of viscoelastic fluids are observed.(5, 45) Alshareedah et al showed for peptide-polyU condensates, a terminal region where G’ and G’’ scaled as expected for a Maxwell fluid and a crossover frequency was observed and indicated the onset of network formation. They observed an increase in viscosity and corresponding increase in terminal relaxation time, as is expected for sticky-Rouse type behavior. To capture faster timescales relevant for biological function, active microrheology performed using optical tweezers can be used to extend the observable frequency range.(3, 4) By measuring the rheological response of PGL-3 and FUS condensates as they age, Jawerth et al ascertained these were aging Maxwell fluids. (3) To our knowledge there have been no measurements of protein based coacervates across the frequency range shown in the present work.

Polyelectrolyte complex coacervates have been characterized over a much broader frequency/time domain than protein-based condensates, largely using time-temperature(46) or time-salt superpositioning,(25, 36, 37, 47) creating master curves that resolve terminal, plateau, transition, and glassy regions. Due to the small range of salt concentrations these protein condensates form over, the sensitivity of protein conformation to salt and temperature, and the diverse range of interactions beyond electrostatic that contribute to the interactome these superpositioning techniques may not be widely applicable to study protein coacervate systems.

The simplicity and ready availability of DLS instruments, combined with the wide frequency range measured makes this approach very well suited to the characterization of biomolecular condensates. Previously, condensate viscosities between ∼0.1 Pa.s and 200 Pa.s have been reported, this range is well accommodated by this technique, plus has the potential to measure much stiffer systems, with moduli up to 10^4^.(48) However, one downside compared to video particle tracking is the loss of spatial information. This means interesting heterogeneity across a sample could be obscured. Another advantage of video particle tracking is that by using protein tracers, the viscosity or viscoelastic response of condensates in cells can be probed directly.(6, 49) By combining DLS with video-particle tracking, this adds another technique to the host of methods currently used to provide detailed insights into condensate properties.

To design functional condensates, it is crucial to understand the underlying interactions that control material properties, including contributions from both disordered and globular domains. In the work presented here we demonstrate that condensates formed between polyK and a model anionic protein with charge differentially distributed between the globular domain and engineered disordered tags. We show that differences in charge distribution result in subtle differences in phase behavior and viscoelasticity. We further show that changes in charge fraction impact viscoelasticity, an effect we attribute to differences composition that influence the number of interactions and how these interactions form a network. Critically, the behavior of condensates containing folded proteins differs significantly from those composed of two linear polypeptides. This work highlights the importance of globular domains when considering condensate properties, showing that the rules for disordered proteins do not necessarily directly translate. Many proteins known to drive phase separation in cellular organelles contain globular domains, their contribution should not be overlooked when considering multivalent interactions and therefore impact on viscoelasticity and other network properties. Looking forward we can begin to design condensates with tailored material properties by engineering both disordered and globular regions, thinking about how domains will behave independently and how they will interact together to control fluid networks.

## Methods

### Protein Expression and Purification

Tag12-GFP plasmid was purchased from Twist and transformed into NiCo21 (DE3) cells, tag6-GFP(-12) and iso-GFP(-12) were previously cloned in the Obermeyer lab and stored as glycerol stocks, plasmids are available from Addgene.

GFPs were expressed in NiCo21 (DE3) cells in LB media. Cells were induced with 1 mM isopropyl β-D-1-thiogalactopyranoside (IPTG) at an OD_600_ of between 0.8-1, grown overnight at 25 °C and harvested by centrifugation (4000 rpm for 20 min). The resulting cell pellet was flash frozen and stored at -80 °C. Cell pellets were defrosted and resuspended in lysis buffer (50 mM sodium phosphate, 300 mM sodium chloride, pH 8) followed by 10 min of sonication (33% duty cycle). Soluble and insoluble fractions were separated by centrifugation at 10,000 xg for 30 min. The resulting supernatant was incubated with Ni-NTA beads equilibrated with Buffer 1 (50 mM sodium phosphate, 300 mM NaCl, and 35 mM imidazole, pH 8.0) for 1 h at 4 °C. Weakly bound protein was removed by washing with Buffer 1. GFP was eluted with Buffer 2 (50 mM sodium phosphate, 300 mM NaCl, and 250 mM imidazole, pH 8). Purified GFP was concentrated by centrifugal ultrafiltration (molecular weight cutoff, MWCO, of 10 kDa) and buffer exchanged into 10 mM Tris (pH 7.4) by dialysis (MWCO of 3.5 kDa). GFP was stored in solution at 4 °C for short term use or mixed with glycerol (10%), frozen in liquid nitrogen, and stored at –80 °C. GFP concentration was determined by measuring absorbance at 488 nm (ε = 83000 M^−1^ cm^−1^) using a Cary 60 UV-Vis spectrophotometer.

### Turbidity Assay

GFP and polycation solutions were combined in tissue culture-treated polystyrene 384-well plates (ThermoFisher) and mixed by pipetting the mixture several times. Turbidity measurements were performed on a Tecan Infinite M200 Pro plate reader. After orbital shaking for 30 s, samples were allowed to sit for 30 s prior to an absorbance measurement at λ = 600 nm. Percent transmittance (%T) was calculated from absorbance (A) using the formula A = 2 − log(%T). Measurements were performed in triplicate across several days.

### Coacervate imaging

Coacervates were prepared in a tissue culture-treated polystyrene 384-well plates treated with a 1% pluronics solution (PLF 127). Samples were imaged on an EVOS™ FL Imaging System equipped with ×60/0.75 numerical aperture (NA) long-working distance Plan fluorite objective. GFP was excited with a 470 +/-22 nm LED. Fluorescence images were acquired with a CCD camera and brightfield with a CMOS camera controlled by EVOS software in manual mode. ImageJ was used to further format and process images. As standard, images were collected 24 h post coacervate formation.

### Particle Tracking Microrheology

Coacervates were prepared in glass bottom 384-well plates (Cellvis) by mixing polyK, 500 nm PEG-functionalized microspheres, and GFP in that order. Carboxy-functionalized microspheres (FluoSpheres, Life Technologies Corporation, 2% solution) were modified with poly(ethylene glycol) (PEG) by addition of an equal volume of PEG-amine (Mr 2 kDa, 5 mg/mL) in MES buffer (50 mM, pH 6) to the microspheres. The solution was incubated for 15 min after which time 1-ethyl-3-(3-dimethylaminopropyl)-carbodiimide (EDAC) was added (final concentration, 25 mM). The pH was adjusted to 6.5 using 1 M NaOH. The solution was incubated overnight. The reaction was then quenched by addition of glycine (final concentration 100 mM). Microspheres were centrifuged at 14,000 g for 15 min, sonicated, and washed with phosphate buffer (50 mM). This was repeated four times, for a total of 5 wash steps.

Imaging was performed using a Nikon Eclipse Ti inverted microscope, equipped with a ×60/1.4 numerical aperture (NA) Plan Apo objective (Nikon) and Perfect Focus System to minimize drift. Red fluorescent beads were excited with the 555 nm line from a LIDA solid-state laser (Lumencor) and collected with a 552 nm edge BrightLine (Semrock) and a FF01 brightline emission filter 572/28 (Semrock). Images were acquired with a sCMOS Andor Zyla VSC-04733 camera controlled using Nikon Elements AR software. 1000 images were collected at 500 ms time intervals. These were exported as tiffs using ImageJ and further analyzed in MatLab using code from the Elbaum-Garfinkle lab, available on online at https://zenodo.org/record/6818910#.YsxnrnbMI2x

From calculated trajectories mean squared displacement (MSD) was calculated as:

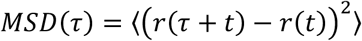

Where r is position, t is time, τ is lag time.

Plotting MSD as a function of time yielded plots with a slope approximately equal to one, indicating that beads were displaying Brownian motion (Figure 3a). This indicates that at the time resolution of these measurements, the coacervates were behaving as Newtonian fluids. Consequently, the diffusion coefficient (D) can be calculated as

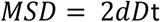

Where d is number of dimensions, in this case 2.

From this, condensate viscosity (η) can then be calculated using the Stokes-Einstein relation:

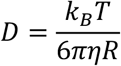

Where *k*_*B*_ is the Boltzman constant, T is temperature, and R is bead radius.

### Dynamic light scattering (DLS)

DLS rheology measurements were performed as described by Cai et al(33) and analysed using their open access method (Version: 0.0.21) which can be found at https://dlsur.readthedocs.io/ and on GitHub at https://github.com/PamCai/DLSuR.

All measurements were performed in a Malvern low-volume quartz cuvette (ZEN2112). Coacervate samples (2 mL) were prepared by adding buffer, polycation, beads, and then GFP in that order directly to the cuvette followed by pipetting to mix. The cuvette was then sealed and left to stand for 30 min, before being placed into a falcon tube and centrifuged at (4000 xg) for 10 min to allow the condensed phase to settle to a layer at the bottom of the cuvette. Measurements were performed in this condensed phase without removal of the dilute phase so as to preserve equilibrium.

Beads that had been surface modified with PEG readily partitioned into GFP condensates. These same surface passivated beads did not readily partition into condensates of polyK-polyD. For example, if beads were injected into the condensates they would migrate to the interface between the dense and dilute phases on a timescale comparable to the DLS measurement length. Unmodified carboxy-beads were consequently used to measure polyK-polyD condensates. Both types of beads were used to measure a condensate with iso-GFP (*f* ^*+*^ = 0.71) and the results were comparable (SI Fig 9).

Autocorrelation curves were collected using a Malvern Zetasizer Nano ZS (633 nm laser) running version 7.12 Zetasizer software and operated in 173° non-invasive backscatter detection mode. The raw intensity autocorrelation function was collected at 4.2 mm, using automatic attenuator selection, for 1000 s. This was followed by fifteen 30 second measurements of the derived photon count rate at a series of positions, used to determine the scattering intensity. Data was exported and MSDs and complex moduli calculated as described in Cai et al(33). Fitting data to extract cross over points and plateau moduli was performed in Matlab.

To perform these measurements, beads were embedded within a large coacervate phase (>12 µL) (SI Fig 7). These beads scatter incident light, the scattering intensity is then measured as a function of time (where *τ* is lag time), and from this the intensity autocorrelation function *g*^2^(*r*) is calculated. Mean squared displacement can be calculated from *g*^2^(*r*) via the intermediate scattering function *g*^1^(τ) where:

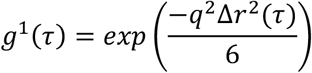

q is the scattering wave vector, calculated as 4πn sin(ϴ/2)/λ where n is refractive index (assumed to be that of water), ϴ is back scatter angle of the detector and λ is wavelength of the laser. The frequency dependent linear viscoelastic shear modulus (*G*\*(*ω*)) can then be calculated as:

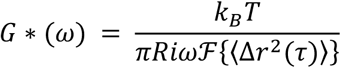

Where ℱ {⟨Δ*r*^2^(*t*)⟩} is the unilateral Fourier transform of mean square displacement. Using this technique, we were able to measure the rheology across a frequency range of 10^−6^ to 10 s in one measurement without the need for time temperature/salt superposition.

Previous condensate rheology results have been fit using a simple Maxwell model(3). The Maxwell model is one of the most fundamental models to describe viscoelastic materials, it describes an exponential decay, with the relaxation time τ where relaxation originates from relaxation of segments between bonds or from motion of whole polymers flowing.

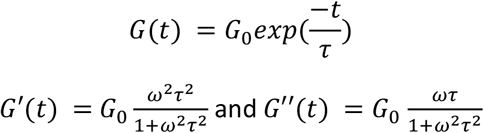

As the timescale for sticky interactions are significantly longer than the timescale for polymer chain relaxation, the stress relaxation can be split into two contributions. A slow component originating from sticker interactions and a fast relaxation due to Rouse motion of the polymer between sticky bonds.

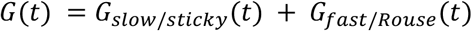

Each of these components can be described by a Maxwell relaxation. Fitting with this two component Maxwell type model we see reasonable agreement at long and intermediate times.

### Dilute Phase Concentration Determination

Dilute phase concentration was determined by first preparing 50 µL coacervate samples in a 384 well plate as described above. Samples were sealed with parafilm and left to settle overnight. Plates were then centrifuged at 4000 xg for 10 min to remove any remaining coacervates from the dilute phase. 40 µL of the dilute phase was carefully pipetted into a new well. Absorbance (A) was measured at 488 nm (ε = 83000 M^−1^ cm^−1^) using a Tecan Infinite M200 Pro plate reader. 40 µL of GFP at a known concentration was used to calculate pathlength (l) and concentrations were calculated using the Beer-Lambert Law, A=cεl. Measurements were performed in triplicate across several days.

