## Supplementary Information for "Viscoelasticity of globular protein-based biomolecular condensates"

**Tuning materials properties of globular protein-based condensates**

**Supplementary Table 1.** Protein sequences

| Protein name | Sequence |
|---|---|
| Iso-GFP | MGHHHHHHGGASKGE**E**LFTGVVPILVELDGDVNGHKFSVRGEGEGDAT**E**GKLTLKFICTTGKLPVPWPTLVTTLTYGVQCFSRYPDHMKQHDFFKSAMPEGYVQERTISFKDDGTYKTRAEVKFEGDTLVNRIELKG**I**DFKEDGNILGHKLEYNFNSHNVYITADKQ**E**NGIKANFKIRHNVEDGSVQLADHYQQNTPIGDGPVLLP**D**NHYLSTQSALSKDP**N**E**D**RDHMVLLEFVTAAGITHGMDE**L**YK |
| Tag6-GFP | MGHHHHHHGGASKGE**E**LFTGVVPILVELDGDVNGHKFSVRGEGEGDAT**N**GKLTLKFICTTGKLPVPWPTLVTTLTYGVQCFSRYPDHMKQHDFFKSAMPEGYVQERTISFKDDGTYKTRAEVKFEGDTLVNRIELKG**I**DFKEDGNILGHKLEYNFNSHNVYITADKQ**K**NGIKANFKIRHNVEDGSVQLADHYQQNTPIGDGPVLLP**D**NHYLSTQSALSKDP**N**E**K**RDHMVLLEFVTAAGITHGMDE**L**YK**DEEEDD** |
| Tag12-GFP | MGHHHHHHGGASKGE**R**LFTGVVPILVELDGDVNGHKFSVRGEGEGDAT**N**GKLTLKFICTTGKLPVPWPTLVTTLTYGVQCFSRYPDHMKQHDFFKSAMPEGYVQERTISFKDDGTYKTRAEVKFEGDTLVNRIELKG**R**DFKEDGNILGHKLEYNFNSHNVYITADKQ**K**NGIKANFKIRHNVEDGSVQLADHYQQNTPIGDGPVLLP**R**NHYLSTQSALSKDP**K**E**K**RDHMVLLEFVTAAGITHGMDE**R**YK**DEEEDDDEEEDD** |

***bold** residues indicate differences between the protein variants

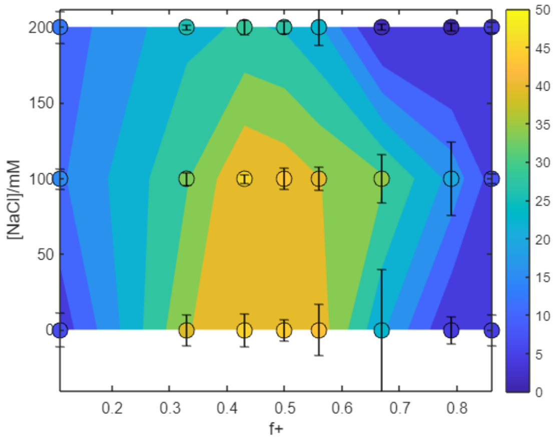


**Supplementary Figure 1**. Turbidity as a function of charge fraction ($f^+$) calculated as $M^+/(M^++M^-)$ where $M^+$ and $M^-$ are charge per polymer and protein respectively measured at 0, 100 and 200 mM NaCl. To prepare coacervates the polyD concentration was held constant at 40 µM in tris (10 mM, pH 7.4) and varying concentration of polyK in tris (10 mM, pH 7.4) was added (5-250 µM).

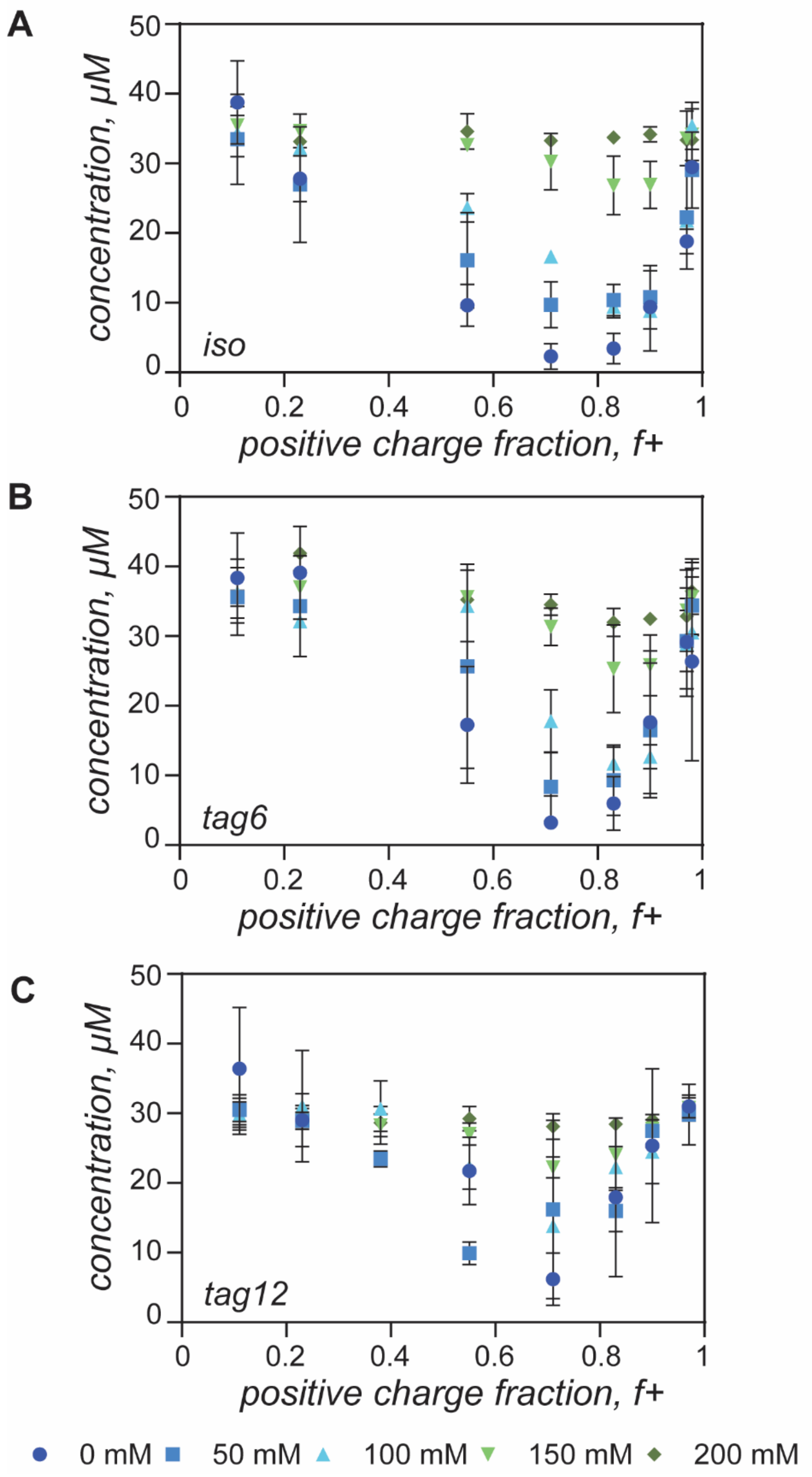


**Supplementary Figure 2.** Concentration of (a) iso-GFP, (b) tag6-GFP, and (c) tag12-GFP in the dilute phase, as measured at 488 nm. GFP (40 µM) with increasing polyK ($f^+$ = 0.11 – 0.98) at 0, 50, 100, 150 and 200 mM NaCl (in 10 mM tris buffer pH 7.4).

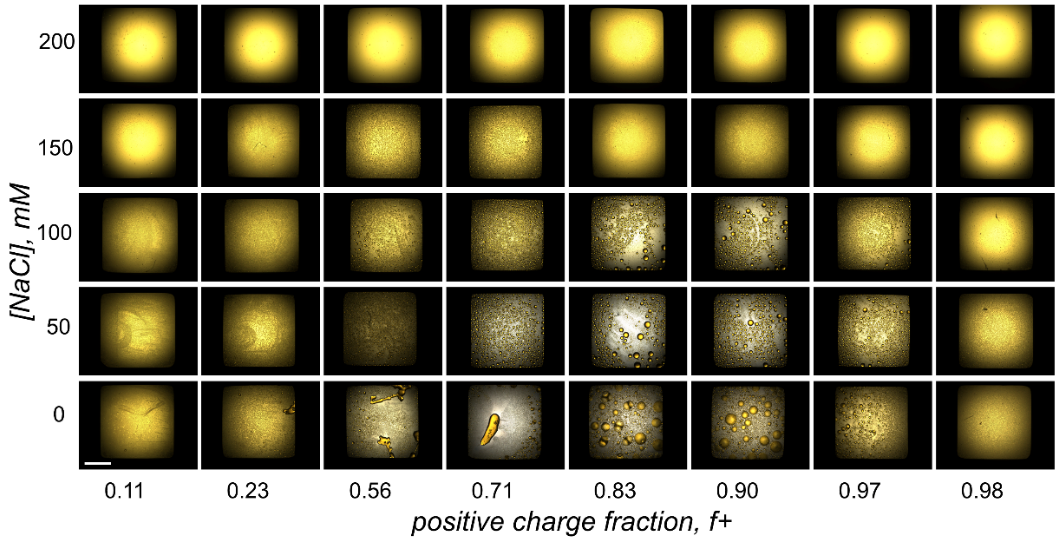


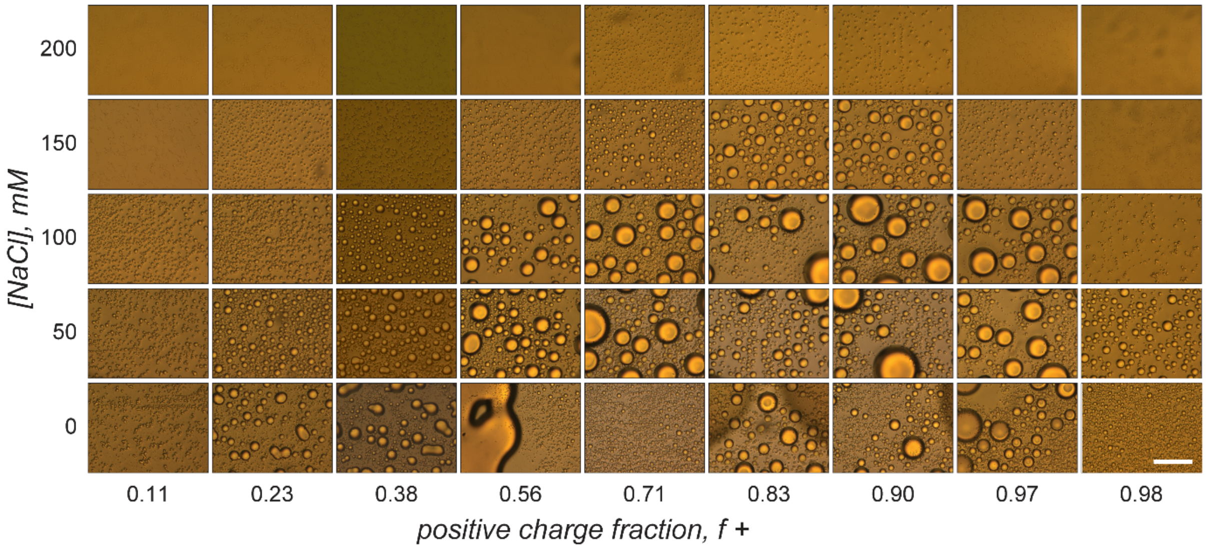


**Supplementary Figure 3.** Brightfield images of iso-GFP (40 µM) with increasing polyK ($f^+$ = 0.11 – 0.98) at 0, 50, 100, 150 and 200 mM NaCl (in 10 mM tris buffer pH 7.4). Top panel is 4x magnification and shows the entire sample, scale bar 1000 µm. Bottom panel is 60x magnification and more clearly shows the difference between liquid and solid-like condensed phases, scale bar 10 µm.

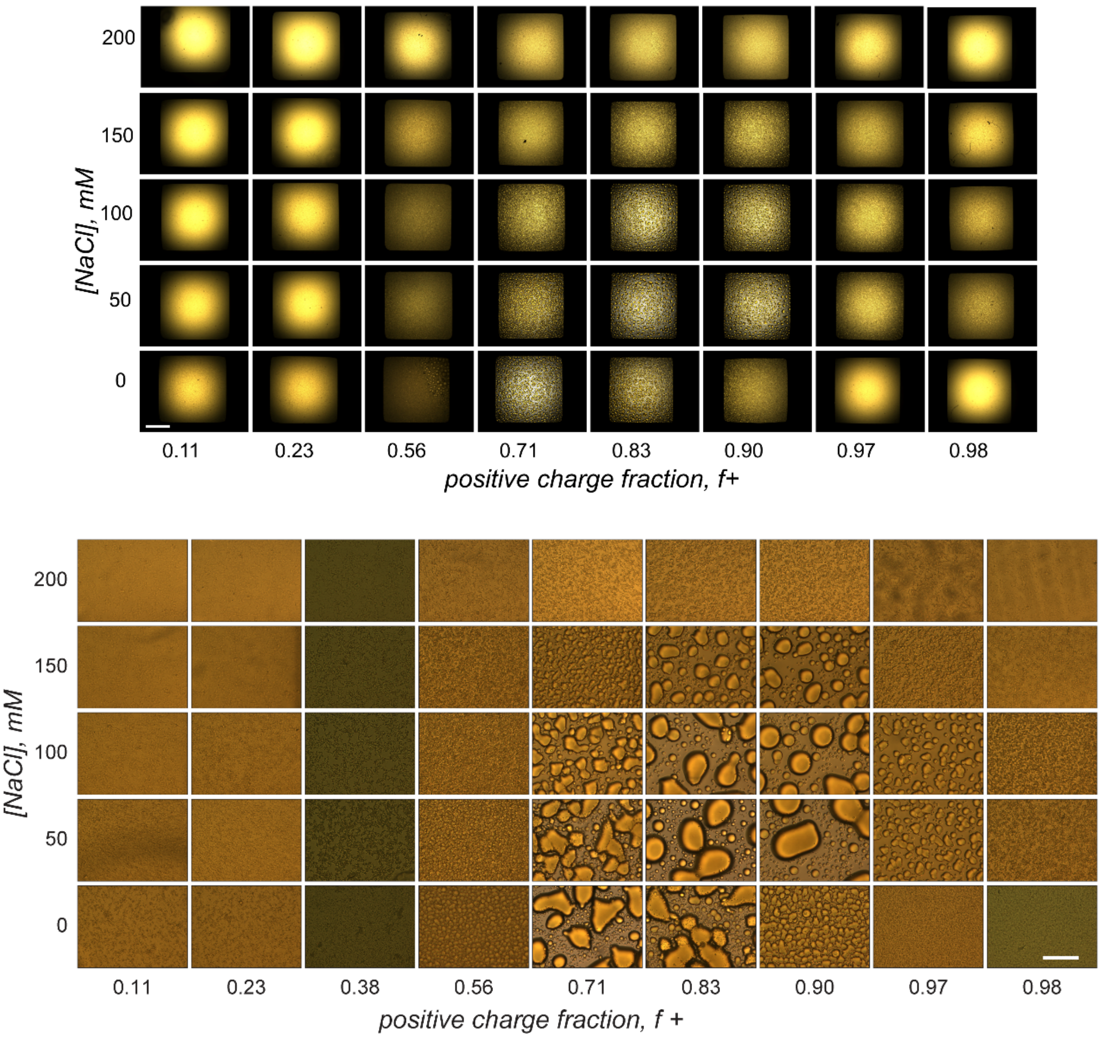


**Supplementary Figure 4.** Brightfield images of tag-6-GFP (40 µM) with increasing polyK ($f^+$ = 0.11 – 0.98) at 0, 50, 100, 150 and 200 mM NaCl (in 10 mM tris buffer pH 7.4). Top panel is 4x magnification and shows the entire sample, scale bar 1000 µm. Bottom panel is 60x magnification and more clearly shows the difference between liquid and solid-like condensed phases, scale bar 10 µm.

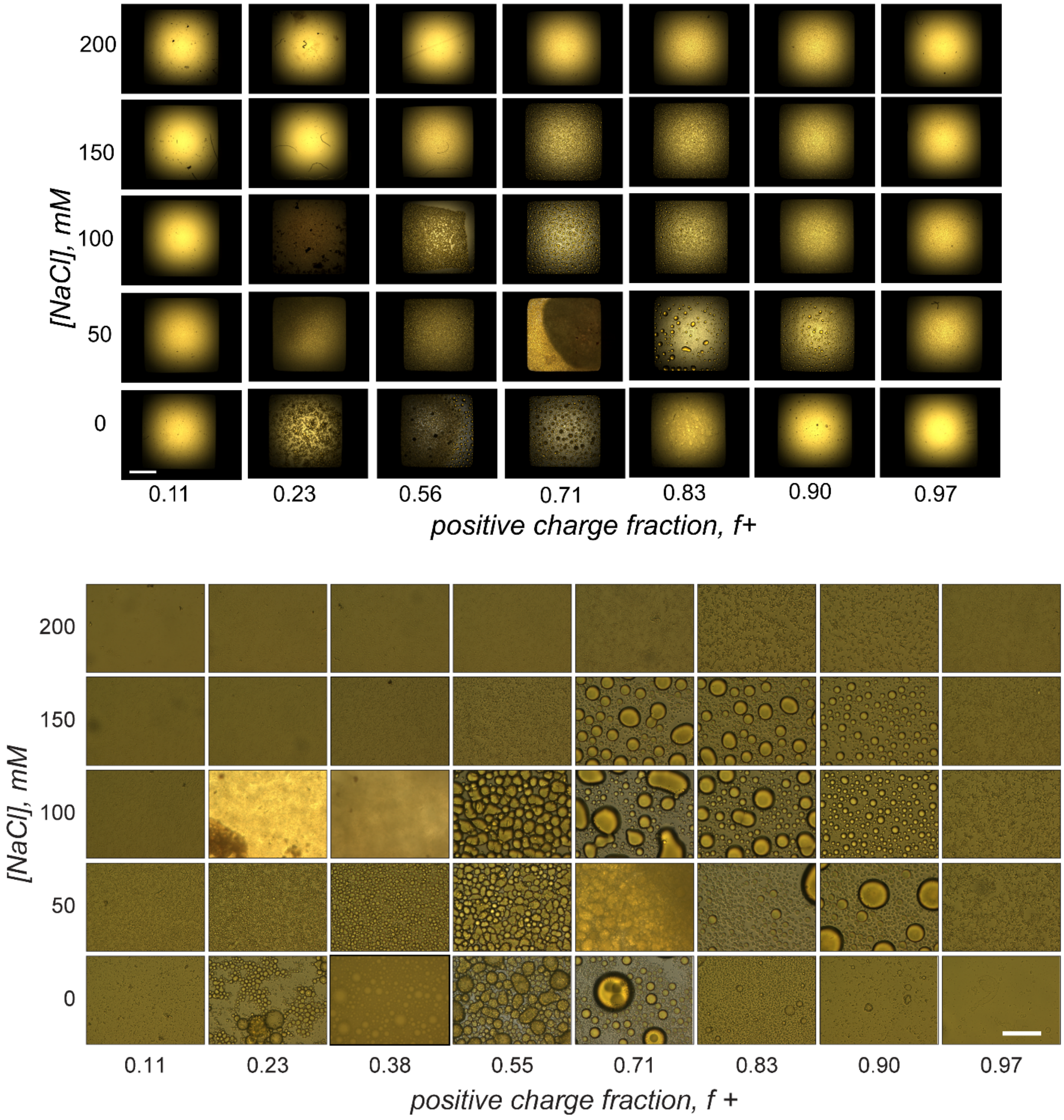


**Supplementary Figure 5.** Brightfield images of tag-12-GFP (40 µM) with increasing polyK ($f^+$ = 0.11 – 0.98) at 0, 50, 100, 150 and 200 mM NaCl (in 10 mM tris buffer pH 7.4). Top panel is 4x magnification and shows the entire sample, scale bar 1000 µm. Bottom panel is 60x magnification and more clearly shows the difference between liquid and solid-like condensed phases, scale bar 10 µm.

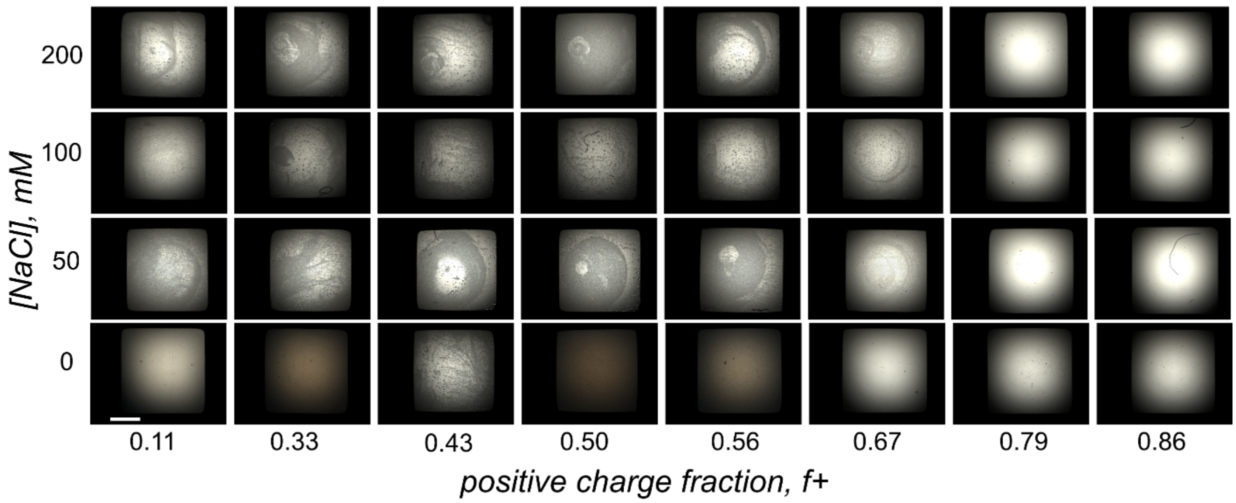


**Supplementary Figure 6.** Brightfield images of polyD (40 µM) with increasing polyK ($f^+$ = 0.11 – 0.86) at 0, 50, 100 and 200 mM NaCl (in tris buffer pH 7.4). The panel shows the entire sample (4x magnification), scale bar 1000 µm.

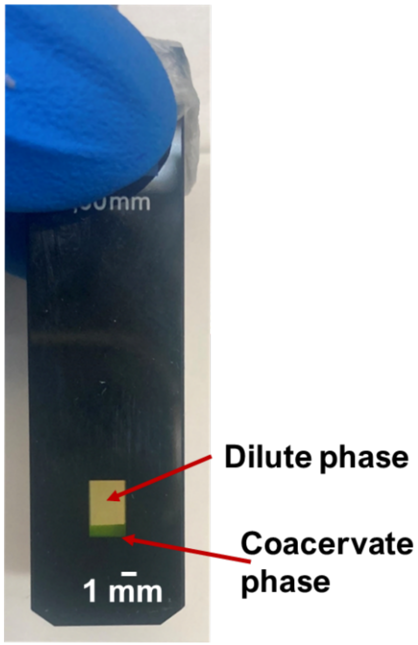


**Supplementary Figure 7.** Quartz micro-cuvette with condensed phase at the bottom of cuvette and dilute phase filling the full 2 mL volume of the cuvette.

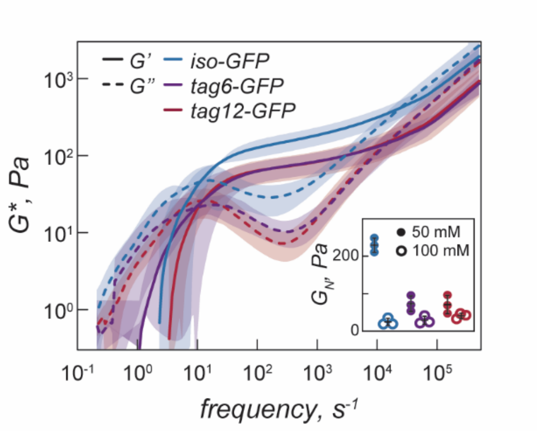


**Supplementary Figure 8.** Complex modulus of iso-GFP (blue), tag6-GFP (purple) and tag12-GFP (red) (40 µM) with polyK (40 µM, f+ = 0.71) at a NaCl concentration of 50 mM. Inset shows plateau modulus ($G_N$) at 50 mM NaCl concentration (solid circles) and 100 mM NaCl (open circles).

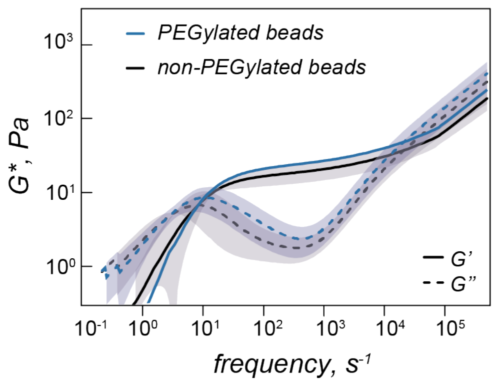


**Supplementary Figure 9.** Complex modulus of iso-GFP with polyK (40 µM, f+ = 0.71) at a NaCl concentration of 100 mM collected using PEGylated 500 nm beads (blue) and non-PEG treated beads (black).

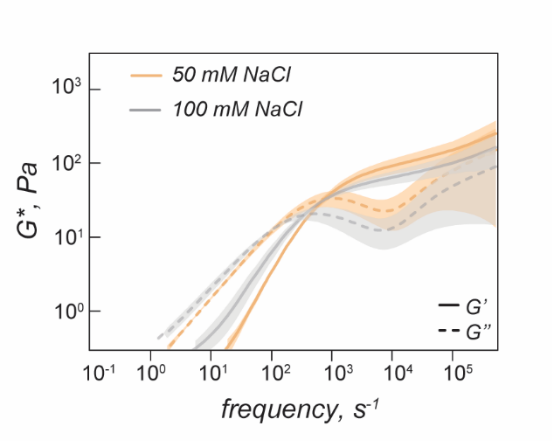


**Supplementary Figure 10.** Complex modulus of polyD30 with polyK (40 µM, f+ = 0.50) at a NaCl concentration of 100 mM (grey) and 50 mM (orange).
